# Genomic Evidence for a Novel Introduction and Intra-host Diversity of DENV-2 in Dar es Salaam, Tanzania

**DOI:** 10.1101/2025.11.17.688789

**Authors:** Silvan Hälg, Frank S.C. Tenywa, Nicole Liechti, Christian Beuret, Sarah J. Moore, Pie Müller

**Affiliations:** Swiss Tropical and Public Health Institute, Allschwil, Switzerland; University of Basel, Basel, Switzerland; Ifakara Health Institute, Bagamoyo, Tanzania; Spiez Laboratory, Spiez, Switzerland; Nelson Mandela African Institute of Science and Technologys, Tengeru, Tanzania

**Keywords:** Aedes aegypti, Arbovirus surveillance, Nanopore sequencing, Phylogenetic analysis, Viral evolution, Genomic epidemiology

## Abstract

Dengue virus (DENV) poses a growing risk in Tanzania, yet its diversity in mosquitoes is poorly understood. Using Nanopore sequencing, we recovered full coding sequences from six DENV-2 positive mosquito pools collected in Dar es Salaam outside outbreak periods. Phylogenetic analysis placed the sequences in a distinct monophyletic clade within genotype II, separate from strains linked to Tanzania’s 2014 outbreak. Instead, they clustered with Asian lineages, showing closest relatedness to DENV-2 from Kenya (2013) and India (2014), with divergence estimated around 2010. Variant profiling identified 212 low-frequency intra-host variants, largely nonsynonymous changes in NS3 and NS5. These results suggest an unrecognized introduction of genotype II now circulating silently in local mosquito populations. Our findings highlight the value of genomic surveillance in vectors for early detection of arboviral threats, even when human cases are absent.

## Introduction

Dengue virus (DENV), a member of the genus *Flavivirus* in the family Flaviviridae, is a significant vector-borne pathogen responsible for a spectrum of clinical outcomes ranging from mild to severe dengue fever (dengue haemorrhagic fever and dengue shock syndrome) (1). It affects approximately 390 million people worldwide annually, with nearly 96 million individuals manifesting clinical symptoms, making it one of the most widespread mosquito-borne viral infections globally (2, 3). The rapid spread of dengue has been fuelled by urbanization, climate change, and increased international travel and trade (2, 4, 5).

Four antigenically distinct serotypes (DENV-1 to −4) circulate in endemic regions like Dar es Salaam, each capable of causing the full clinical disease spectrum (6). While primary infection provides long-term immunity against that specific serotype, heterologous secondary infections can lead to more severe disease due to antibody-dependent enhancement (7, 8). Each serotype is further divided into multiple genotypes (genotype I-V for DENV-1 and −2, and I-VI for DENV-3 and −4) with distinct geographic distributions and epidemiological patterns (9).

Molecular epidemiology, particularly whole-genome sequencing, has become essential for tracing DENV introduction, lineage replacement, and viral adaptation to vectors and hosts (10, 11). Phylogenetic studies have highlighted increasing instances of intercontinental virus movement, with rising reports of Asian DENV genotypes in Africa (12, 13).

Despite this, Africa, and Tanzania in particular, remains underrepresented in genomic surveillance data. Tanzania has witnessed multiple dengue outbreaks since 2010, including major episodes in 2014 (predominantly DENV-2, genotype II, Cosmopolitan) and 2019 (DENV-1) (12). These outbreaks have been linked phylogenetically to Asian countries such as China and Singapore (12). However, few studies have sequenced viruses directly from vectors, and information on viral evolution across mosquito reservoirs remains limited.

Additionally, there is growing evidence that intra-host viral diversity, especially within non-structural genes (NS) like NS3 and NS5, plays a vital role in dengue’s adaptive evolution. Studies in human and mosquito hosts have reported selection pressures on NS genes during natural DENV infection (14). Intra-host diversity has been linked to viral replication fitness and quasi-species dynamics, emphasizing its relevance for surveillance and prediction of transmission risk (15).

In a previous longitudinal xenomonitoring study, DENV-2 has been detected in *Aedes aegypti* mosquitoes across Dar es Salaam during inter-epidemic period, indicating silent, persistent urban transmission (16). Such low-level transmission without detected cases is increasingly recognized as a crucial factor in maintaining dengue virus circulation and potentially seeding future epidemics (17, 18). To better understand the evolutionary origins, genomic diversity, and potential introduction routes of these mosquito-derived DENV-2 strains, we applied Oxford Nanopore sequencing to generate full-genomes from six DENV-2–positive mosquito pools.

This study represents the first detailed genomic and phylogenetic characterization of DENV-2 circulating within mosquito vectors in Dar es Salaam, Tanzania. We integrate whole-genome and gene-specific phylogenies with intra- and inter-sample variant analyses to determine whether these viruses represent ongoing local lineages or newly introduced genotypes. Our findings yield critical data for enhancing genomic surveillance strategies and inform public health authorities in Tanzania and the broader East African region.

## Methods

### Sample origin and RNA extraction

A total of six DENV-2 positive mosquito samples were used for sequencing. Five of these viruses were identified via reverse transcription quantitative PCR (RT-qPCR) during a two-year longitudinal study conducted in Dar es Salaam, Tanzania, as previously described (16). The sixth DENV-2-positive sample was detected in a pool of *Ae. aegypti* mosquitoes collected in 2022 from the same study area but processed independently.

For this latter sample, 600 µl of InhibitEx Buffer (Qiagen AG, Hilden, Germany) was added to a pool of 10 mosquitoes. The sample was homogenized twice for 30 seconds using a QIAGEN TissueLyser II, then centrifuged at 3,220 × g for 5 min. A 100 µl aliquot of the supernatant was combined with 400 µl AVL buffer (Qiagen). Nucleic acid extraction was performed using the DNA and Viral NA Large Volume Kit (Roche Diagnostics, Rotkreuz, Switzerland) on the MagNa Pure 96 System (Roche Diagnostics). The remaining five DENV-2-positive samples were processed and extracted following the methods described in the original study (16).

### cDNA synthesis and amplicon generation

Extracted RNA was reverse-transcribed using SuperScript IV Reverse Transcriptase (Thermo Fisher Scientific, Waltham, Massachusetts, U.S.A.) with random hexamers. Full genome amplification of DENV-2 was performed using a tiling multiplex PCR approach. Primer sequences were used as published for DENV-2 in the protocol by Su et al. (19), which enables serotype-specific and multiplexed amplification of dengue virus genomes directly from clinical or entomological samples. The primers generate ∼400 bp overlapping amplicons covering the entire coding region of the genome.

### Library preparation and Nanopore sequencing

PCR products were purified using AMPure XP beads (Bioconcept AG, Allschwil, Switzerland). Purified amplicons were barcoded using the Native Barcoding Expansion Kit (Oxford Nanopore Technologies, Oxford, U.K.) and prepared for sequencing with the Ligation Sequencing Kit (SQK-LSK114) (Oxford Nanopore Technologies, Oxford, U.K.). Libraries were loaded onto a MinION R10.4.1 flow cell and sequenced on the GridION platform (Oxford Nanopore Technologies, Oxford, U.K.) using MinKNOW software for live base-calling (Guppy v6.4.2, high-accuracy mode), real-time demultiplexing and quality filtering (Q>10).

### Genome assembly and consensus generation

Primer sequences from raw reads were trimmed using *cutadapt* (20). Reads were filtered for quality >20 with *Filtlong* (21) and then mapped to a reference DENV-2 genome (GenBank: MT982148.1) using *Minimap2* v2.28-r1209 (22) and indexed with *SAMtools* (23). Consensus sequences and variant calling were performed using *iVar* v1.4.3 (24). Positions with coverage below 10× were masked with “N” to ensure high-quality consensus sequences. Complete genome sequences were submitted to GenBank under accession numbers PV834971–PV834976.

### Multiple sequence alignment and phylogenetic analysis

To determine the genotype of DENV-2 sequences and contextualize them within the global DENV-2 landscape, three separate phylogenetic analyses were performed: serotype identification, complete genome-based phylogeny, and E protein-based phylogeny. All analyses used *MAFFT* v7 (25) for sequence alignment and *BEAST* v2.7.7 (26) for tree reconstruction.

To determine the genotype of the sequenced DENV-2 isolates, consensus genomes from the six samples were aligned against representative DENV-2 reference sequences encompassing all six recognized genotypes (Genotypes I–VI). Tip dates were not included in the genotype analysis, as temporal information is not required for establishing genotype identity based on phylogenetic clustering. A generalized time reversible (GTR) substitution model with gamma-distributed rate variation across four categories, a relaxed log-normal molecular clock with a mean clock rate of 0.001 (27), and a Bayesian Skyline coalescent tree prior were applied. XML input files were generated using *BEAUti*, part of the BEAST suite. The Markov Chain Monte Carlo (MCMC) was run for 100 million generations to ensure adequate sampling of the posterior distribution. Convergence and effective sample size were assessed using Tracer v1.7.2 [36], with a 10% burn-in applied during maximum clade credibility (MCC) tree generation using *TreeAnnotator* within *BEAST*. Final trees were visualized in *FigTree* v1.4.4 (28).

To place our samples into the broader phylogenetic context of DENV-2 genotype II, a total of 336 unique complete DENV-2 genotye II genomes were retrieved from GenBank using Biopython (29). These, along with our consensus genomes, were aligned with MAFFT v7 using default settings. For this analysis, tip dates were incorporated, and phylogenetic inference was performed in BEAST using a GTR + gamma (4 categories) substitution model, a relaxed log-normal molecular clock with a mean clock rate of 0.001 as a starting value (27), and a Bayesian Skyline coalescent prior. The MCMC was run for 100 million generations, and the MCC tree was produced after removing 10% burn-in. Importantly, the final MCC tree was rooted to a historical DENV-2 genotype II reference sequence from 1944 (GenBank accession: KM204118.1) to contextualize temporal divergence. The tree was visualized in *FigTree* (28).

To increase phylogenetic resolution within the E gene, we conducted a separate analysis including 1,132 unique DENV-2 genotype II E gene sequences, derived from GenBank, full genomes, and our six newly generated sequences. The E protein is commonly used for phylogenetic analysis because it plays a critical role in viral entry and host immune response, leading to high genetic variability that reflects the virus’ evolutionary dynamics. As a result, it provides strong discriminatory power for differentiating genotypes and tracking viral transmission. Due to its importance, many E gene sequences are publicly available, making it a practical and informative target for molecular epidemiology. Detailed methods and the resulting phylogenetic tree are provided in the Supplementary File.

### Variant analysis

To characterize within-sample viral population a consensus-based alignment was employed. Sequencing reads were aligned using *Minimap2* v2.24 (22). Alignment outputs were processed using *Samtools* v1.14 (23) for file format conversion, sorting, and indexing prior to downstream variant calling.

To reduce mapping bias and improve variant detection sensitivity, sequencing reads from each mosquito pool were aligned to a sample-specific consensus genome, generated from the same dataset. This approach improves resolution of low-frequency variants, which may reflect quasispecies diversity or multiple infections within the mosquito pool, by minimizing false positives due to divergence from a distant reference genome. By aligning reads to their own consensus, we reduce the influence of fixed inter-sample mutations and improve variant detection in highly polymorphic regions.

Variants were called using the *iVar* pipeline (24). Primer sequences were trimmed using *cutadapt* (20). Variant calling was performed using *iVar* variants, with a minimum base quality threshold of 20. Only variants marked with the “PASS” flag were retained for further analysis. Additional filtering was applied as follows: p-value < 0.05, total read depth > 400, alternate allele frequency > 5%, alternate base must be one of A, T, G, or C (ambiguous calls such as -n, +n, or indels were excluded). Stop codons were also excluded from analysis. Variants were classified as synonymous or nonsynonymous by comparing the reference and alternate amino acids. Codon positions were translated into global polyprotein coordinates based on gene annotations to facilitate gene-level comparisons. Processed variant data were analyzed and visualized in R, using the following packages: *ggplot2* (30), *tidyverse* (31), and *ComplexUpset* (32, 33) for generating the upset plot. A custom genome schematic of the DENV-2 polyprotein, showing the structural (Capside (C), precursor membrane protein gene (prM), envelope gene (E)) and non-structural (NS1–NS5) coding regions, was generated and overlaid beneath variant plots to provide visual and functional context for mutation distribution.

## Results

### Nanopore sequencing

Using Nanopore sequencing, full genomes (>10,000 nt) were successfully recovered from all six DENV-2-positive mosquito pools and yielded between 260,000 to 5,000,000 reads per sample, with a genome coverage between 6,600 to 135,000-fold.

### Phylogenetic analysis

To determine the genotypes of the newly sequenced DENV-2 samples from Dar es Salaam, a phylogenetic tree was constructed using a reference panel representing the known diversity of DENV-2 genotypes. Our samples were clustered within the genotype II clade, confirming their genotype assignment (Figure 1). This initial analysis provided a robust framework for detailed downstream evolutionary and epidemiological analyses.

**Figure 1:**
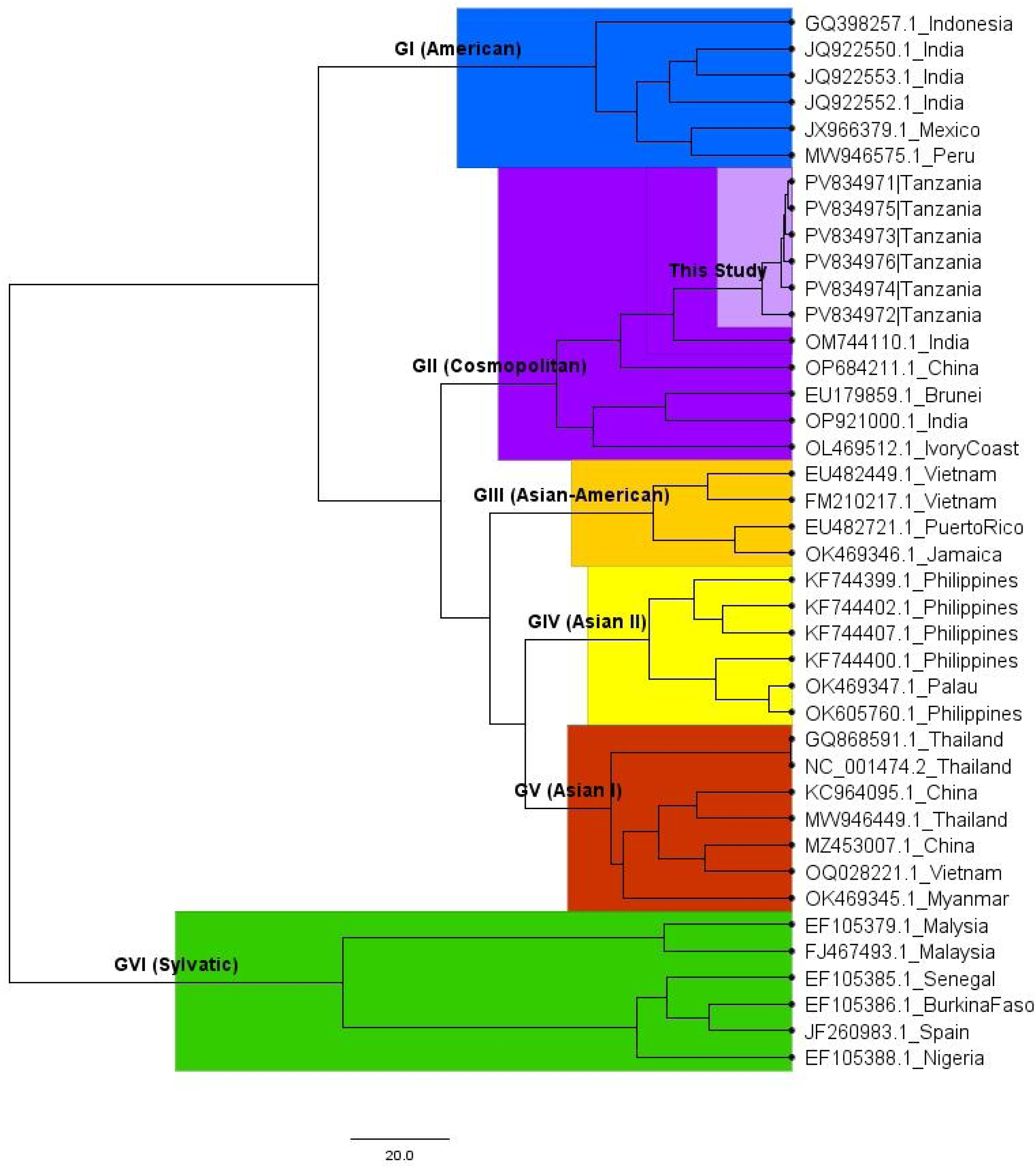
Phylogenetic tree of DENV-2 all genotypes. The tree was inferred using a GTR nucleotide substitution model with gamma-distributed rate variation across sites, discretised into four categories (+G4). A relaxed log-normal molecular clock model was applied with a mean clock rate of 0.001 substitutions/site/year. The tree prior followed a coalescent Bayesian Skyline model to account for demographic changes over time. The Markov Chain Monte Carlo (MCMC) chain was run for 100 million iterations to ensure convergence and adequate sampling of the posterior distribution.

Next, the analysis was expanded by extending the data set with 336 unique complete DENV-2 genotype II genomes that had been sampled globally. Using tip-dated sequences and a relaxed molecular clock model, a time-scaled phylogeny was inferred to investigate the temporal relationship and evolutionary history within this genotype. The estimated mean substitution rate for the DENV-2 genotype II was 9.17 × 10⁻⁴ substitutions/site/year (95% Highest Posterior Density interval, HPD: 8.24 × 10⁻⁴ – 1.01 × 10⁻³).

All Tanzanian DENV-2 sequences from this study clustered on a single terminal branch but did not group with previously reported complete DENV-2 genomes from Tanzania collected in 2014. Phylogenetic analysis suggests these sequences began diverging as early as 1964 (95% highest posterior density [HPD]: 1954–1976) (see Figure 2). Instead, the newly characterized isolates demonstrate closer phylogenetic relatedness to strains from Kenya (isolated in 2013) and India (isolated in 2014). Molecular clock analyses suggest that the divergence between the new Tanzanian and Kenyan isolates occurred around 2011 (95% HPD: 2011–2012), with both lineages further diverging from the Indian isolate around 2010 (95% HPD: 2009–2011). A more ancestral node connects these to a Singaporean isolate, with an estimated divergence around 1999 (95% HPD: 1994–2003) (Figure 2).

**Figure 2:**
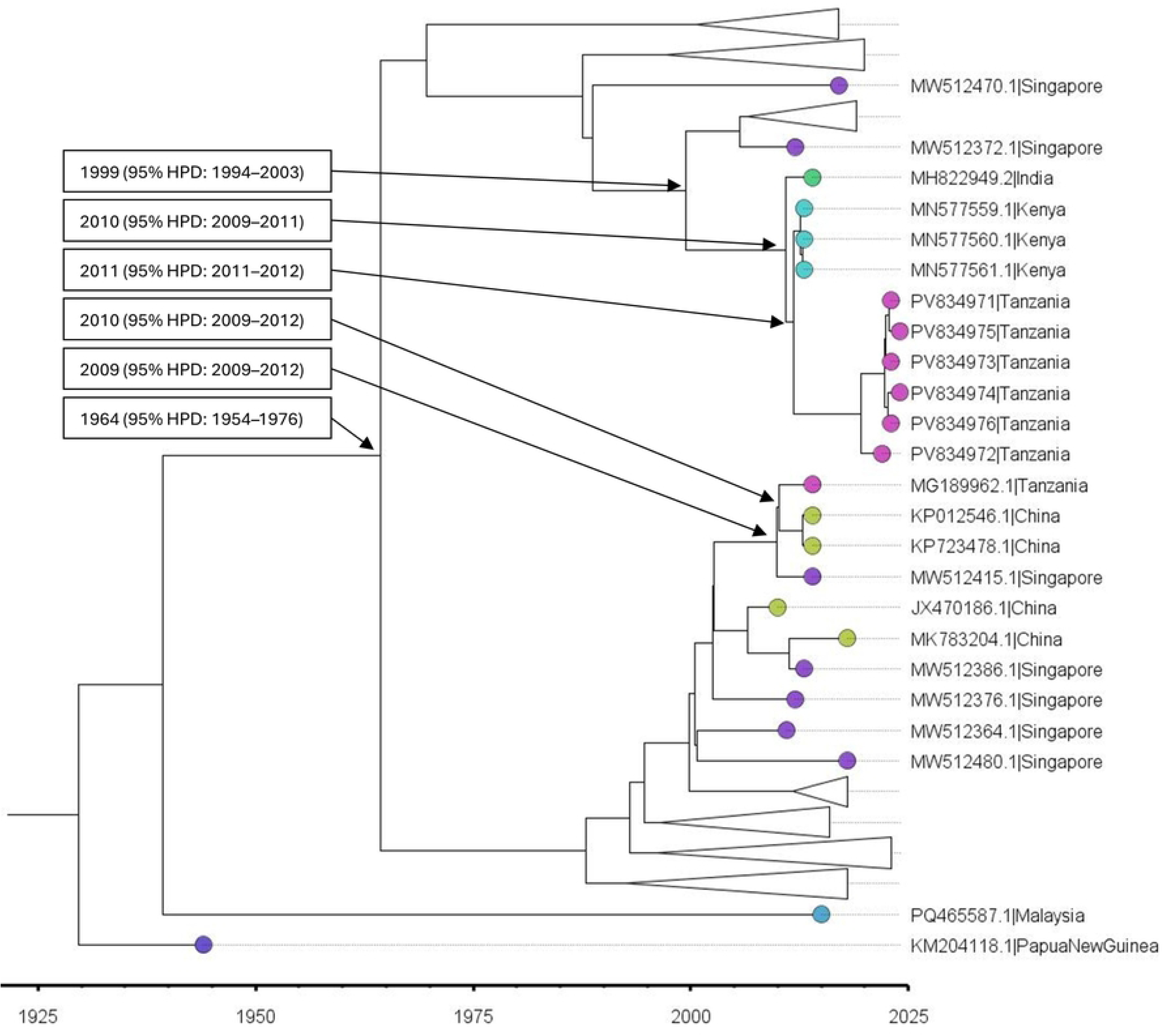
Phylogenetic tree of DENV-2 genotype II complete genome sequences. The tree was inferred using a GTR nucleotide substitution model with gamma-distributed rate variation across sites, discretised into four categories (+G4). A relaxed uncorrelated log-normal molecular clock model was applied, with the mean clock rate fixed at 0.001 substitutions/site/year. To account for demographic changes over time, a coalescent Bayesian Skyline tree prior was employed. The Markov Chain Monte Carlo (MCMC) analysis was run for 100 million iterations, sampling every 10,000 steps, with the first 10% discarded as burn-in. Convergence was confirmed by effective sample size (ESS) values exceeding 200 for all parameters. Posterior probabilities and 95% highest posterior density (HPD) intervals support divergence time estimates. Tips are coloured by country of origin to illustrate the geographic distribution of sequences.

A parallel analysis based on the E gene dataset yielded consistent topological patterns and divergence estimates, further supporting these findings (Supplementary Figure 1).

### Variant analysis

Using a consensus alignment strategy, which involved realigning sequencing reads to each sample’s own consensus genome, 212 intra-host variants were identified across the six samples, with 170 being private to individual samples (Figure 3). Within individual samples, most variants exhibited allele frequencies between 5% and 20%, consistent with low-frequency variants, likely representing intra-host viral subpopulations (Supplementary Figures 2-7).

**Figure 3:**
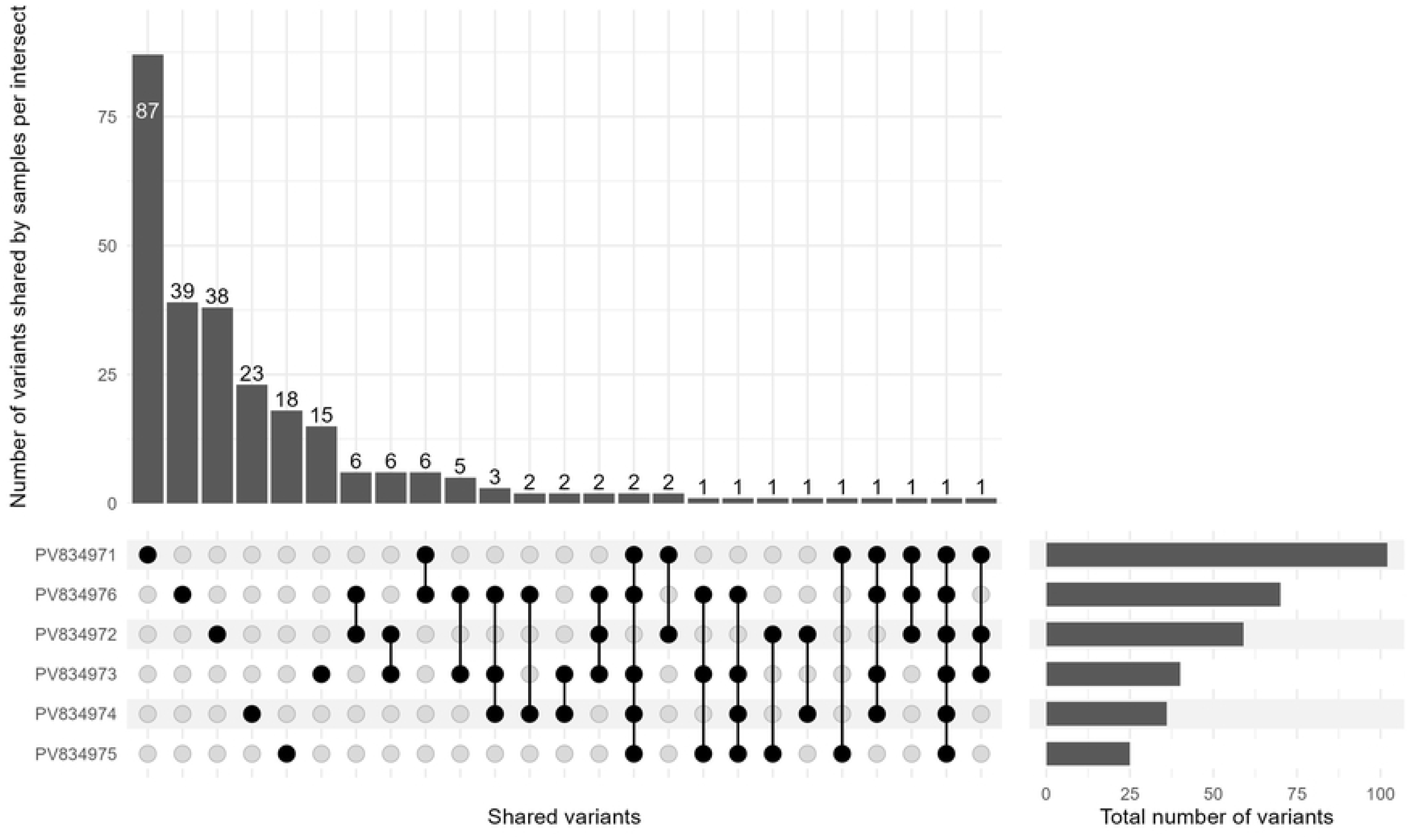
Distribution of shared mutations among samples. Each bar in the intersection matrix (top panel) represents the number of variants shared by the corresponding combination of samples a indicated by the black dots in the bottom matrix. Single-sample variants are shown as individual bars, while multi-sample intersections highlight variants common to two or more samples. The plot illustrates both unique and overlapping variant profiles, allowing comparison of variant sharing across the cohort.

Nonsynonymous mutations were broadly distributed throughout the polyprotein and outnumbered the synonymous mutations in most samples. They were particularly enriched in non-structural genes, especially NS3, NS4B and NS5 (Figure 4 and Supplementary Figures 2 to 7), indicating a potential enrichment of functionally relevant variation at the intra-host level.

**Figure 4:**
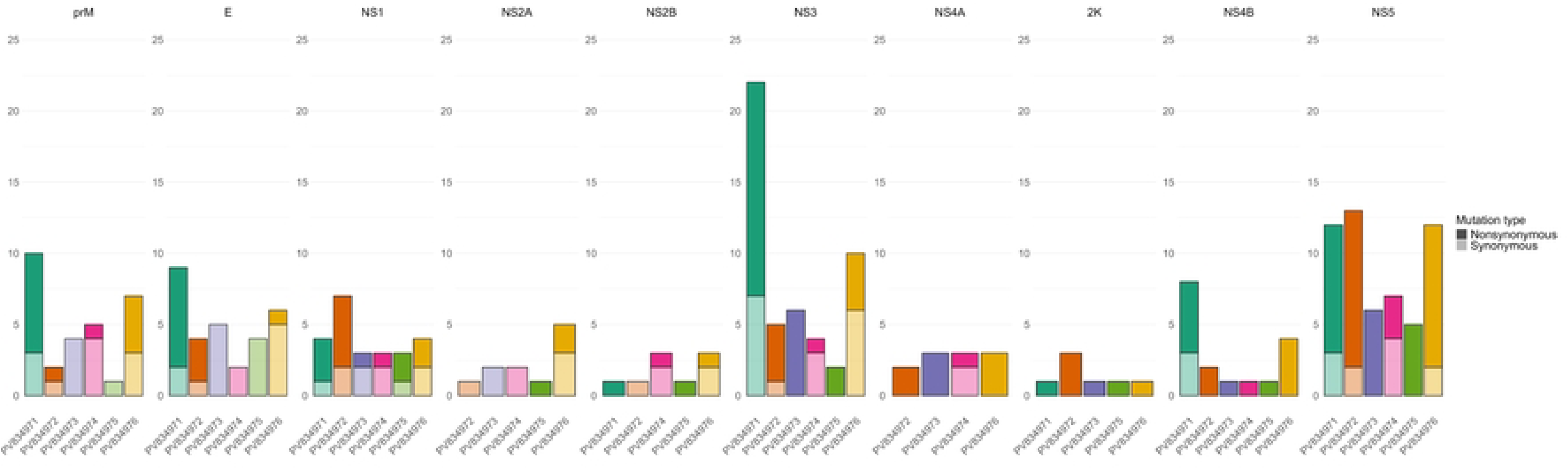
Distribution of synonymous and nonsynonymous mutations across the six DENV-2-positive *Aedes aegypti* pools. Stacked bar plots display the number of mutations detected for each gene of the six DENV-2 samples. Each panel corresponds to a distinct viral gene, arranged in genomic order. Colours represent individual samples, with light colours representing synonymous and dark colours showing nonsynonymous mutations. No mutations were identified in the capsid region therefore it is not shown.

## Discussion

This study reports the detection of DENV-2 genotype II in *Ae. aegypti* mosquitoes collected between outbreaks in Dar es Salaam, Tanzania. Further analysis revealed a complex landscape of low-frequency genetic diversity within the mosquito-derived DENV-2 populations with several predominating nonsynonymous mutations in the NS3, NS4B, and NS5 coding regions.

Comparative genomic and phylogenetic analyses revealed that the newly identified Tanzanian DENV-2 genotype II isolates form a distinct cluster from those linked to the 2014 outbreak (34). The new isolates cluster with Kenyan (2023) and Indian (2014) ones, supporting the hypothesis of multiple temporally staggered introductions into the region (35). Molecular clock analyses indicate that the Tanzanian DENV-2 sequences reported here share a most recent common ancestor with Asian lineage viruses around 2009–2011. This divergence predates the lineage responsible for the 2014 outbreak in Tanzania, which clusters instead with viruses from China and Singapore sampled in 2014. Our sequences therefore represent a distinct introduction, separate from the 2014 outbreak lineage (13). The most recent common ancestor of the two lineages found in Tanzania date back to around 1964. The deep divergence between the newly detected isolates and the 2014 outbreak strains further supports the hypothesis of independent introductions with distinct epidemiological histories, echoing findings from broader regional arbovirus surveillance efforts (2).

A similar virus flow has previously been documented between East Africa and South Asia (12), and recent studies from Kenya and Djibouti have documented diverse DENV-2 lineages, some also closely related to Asian strains (12, 36–38). Altogether, they emphasize that East Africa serves as a convergence point for interregional dengue virus traffic with complex and dynamic transmission patterns in the region (39). The observation of at least two independent introductions of DENV-2 genotype II into Tanzania parallels patterns seen in other hyperendemic regions such as Brazil and Southeast Asia, where multiple co-circulating lineages within a single serotype are common (40, 41). These data support the notion that long-range viral movement, likely facilitated by international travel and trade, is a major driver of dengue virus genetic diversity in Africa.

The predominance of nonsynonymous over synonymous variants suggests selection for specific traits, such as evasion of host immune responses or adaptations to tissue-specific environments within the mosquito vector (14, 42). In the present study, nonsynonymous variants were particularly frequent within sequences coding for non-structural proteins NS3, NS4B, and NS5. This observation aligns with prior evidence that these regions are hotspots for functional adaptations in flaviviruses (14, 42, 43). Notably, NS5, which encodes the RNA-dependent RNA polymerase, is known to undergo adaptive changes influencing viral replication fidelity and immune evasion (44). The observed intra-host evolutionary pattern aligns with results from studies on global RNA virus evolution (14, 42). They found that increased intra-host diversity selects for fitter variants that eventually lead to outbreaks. Indeed, viral evolution within vectors is a critical, yet often underappreciated, component of arboviral epidemiology (44–47).

The findings from the present study emphasize the importance of continuous molecular surveillance to detect new introductions before they trigger outbreaks. The genetic divergence between currently circulating isolates and those from 2014 may also have implications for diagnostic accuracy and local immunity, particularly given the higher rates of severe disease associated with secondary DENV-2 infections (48, 49).

While this study provides valuable insights into the genomic diversity and multiple introductions of DENV-2 in Dar es Salaam, several limitations should be acknowledged. Although screening was performed for all four DENV serotypes throughout the study period, only six DENV-2 genotype II samples were detected and sequenced. This limited number of samples restricts the depth of viral diversity captured and may not fully represent the broader circulating viral population. The analysis was based on viral RNA extracted from pooled mosquito samples rather than from individual mosquitoes or human cases, potentially obscuring more detailed intra-host diversity and transmission dynamics. Furthermore, the relatively small number of genomes restricts our ability to comprehensively capture evolutionary patterns within the region. Temporal sampling gaps and the absence of contemporaneous clinical sequences further constrain our capacity to directly link vector-borne viral diversity with human infections. Future research should prioritise increasing sample sizes, incorporating sequencing from individual mosquitoes and human cases, and integrating epidemiological data to better elucidate transmission networks and viral evolution. Longitudinal studies combining vector and human surveillance will be essential to clarify drivers of viral persistence and emergence. Additionally, functional characterisation of nonsynonymous mutations, particularly within NS3, NS4B, and NS5, will enhance understanding of their effects on viral fitness, vector competence, and immune evasion.

In conclusion, this study underscores the importance of entomological genomic surveillance for early detection of emerging arboviral threats in the absence of reported human cases. Dengue control requires integration of human case monitoring with vector-based genomic surveillance. Such integrated approaches have been instrumental in revealing mechanisms of viral persistence, adaptation, and spread, thereby informing more targeted control strategies against both the virus and its mosquito vectors [53, 54]. Our findings underscore the importance of sustained genomic surveillance particularly in Tanzania, given its complex and evolving transmission landscape to better anticipate and mitigate potential future outbreaks.

## Author Contributions

Conceptualisation: S.H. and P.M.; formal analysis: S.H.; investigation: S.H., F.S.C.T., N.L.; methodology: S.H., N.L.; supervision; P.M.; writing original draft: S.H.; visualisation: S.H.; writing review and editing: N.L., C.B., S.J. M., P.M. and F.S.C.T; funding acquisition: C.B., P.M.; All authors have read and agreed to the published version of the manuscript.

## Informed Consent Statement

Not applicable.

## Data Availability Statement

The collection and use of mosquito specimens in this study were conducted in compliance with the Nagoya Protocol on Access and Benefit-Sharing. Sampling and export of mosquito material were authorized by the Tanzanian authorities under research permit No. #IHI/CED/DSM/2022/476, issued by the Vice President’s Office. The subsequent transport of specimens to Switzerland for viral metagenomic analysis was carried out in accordance with the terms of this permit and applicable international regulations.

Complete genomes from the six dengue viruses are available on GenBank under accession numbers PV834971, PV834972, PV834973, PV834974, PV834975 and PV834976.

## Acknowledgments

The authors would like to thank Ifakara Health Institute staff for their support. We would like to thank Haji Makame, Osward Dogan and Harubu I. Mapipi for the collection of the mosquito. ChatGTP was used to improve text readability.

## Appendices

Supplementary File

## Abbreviations

C: Capside gene
DENV: Dengue virus
E: Envelope gene
GTR: Generalized time reversible
HPD: Highest Posterior Density
MCC: maximum clade credibility
MCMC: Markov Chain Monte Carlo
NS: Non-structural gene
ONT: Oxford Nanopore Technology
prM: Precursor membrane protein gene

